# Bayesian inference of distributed time delay in transcriptional and translational regulation

**DOI:** 10.1101/608596

**Authors:** Boseung Choi, Yu-Yu Cheng, Selahittin Cinar, William Ott, Matthew R. Bennett, Krešimir Josić, Jae Kyoung Kim

## Abstract

**Motivation:** Advances in experimental and imaging techniques have allowed for unprecedented insights into the dynamical processes within individual cells. However, many facets of intracellular dynamics remain hidden, or can be measured only indirectly. This makes it challenging to reconstruct the regulatory networks that govern the biochemical processes underlying various cell functions. Current estimation techniques for inferring reaction rates frequently rely on marginalization over unobserved processes and states. Even in simple systems this approach can be computationally challenging, and can lead to large uncertainties and lack of robustness in parameter estimates. Therefore we will require alternative approaches to efficiently uncover the interactions in complex biochemical networks.

**Results:** We propose a Bayesian inference framework based on replacing uninteresting or unobserved reactions with time delays. Although the resulting models are non-Markovian, recent results on stochastic systems with random delays allow us to rigorously obtain expressions for the likelihoods of model parameters. In turn, this allows us to extend MCMC methods to efficiently estimate reaction rates, and delay distribution parameters, from single-cell assays. We illustrate the advantages, and potential pitfalls, of the approach using a birth-death model with both synthetic and experimental data, and show that we can robustly infer model parameters using a relatively small number of measurements. We demonstrate how to do so even when only the relative molecule count within the cell is measured, as in the case of fluorescence microscopy.

## 1 Introduction

The dynamics of intracellular processes are determined by the structure and rates of interactions between different molecular species. However, stochasticity and limitations of experimental methods make it difficult to infer the characteristics of these interactions from data. On the single-cell level, different molecular species can occur in small number, correlate with phenotype, and localize within different parts of the cell. The resulting dynamics can thus be highly variable over time, and across the population. Averaging over such fluctuations can lead to inaccurate representations of the underlying biology [1], and inference methods therefore need to account for stochasticity within individual cells, and variability across the population [2–5].

Different statistical approaches have been developed to fit stochastic models to data from single-cell assays, offering a window into the dynamical processes within individual cells [6–13]. Among these, Bayesian methods have been particularly promising. To apply Bayesian techniques, one typically assumes a model for the network of interactions, postulates a prior over model parameters, and uses experimental data to determine a posterior and estimates of unknown parameters [6, 13–15]. However, Bayesian approaches can suffer from the curse of dimensionality [16], and are thus difficult to implement directly when the number of parameters is high, or the network of interactions is large. The problem is exacerbated when the system is not fully observed, as here one must marginalize over the unobserved components of the system.

One way to circumvent this problem is to replace uninteresting or unobserved reactions with time delays [17–21]. For instance, the production of regulator proteins requires on the order of minutes: Production involves transcription, translation, and post-translational steps such as protein folding, oligomerization and maturation [2, 22]. Rather than model each step individually [23, 24], one can describe protein production by an effective, random delay that represents a sequence of noisy biochemical processes with fluctuating completion times [25]. Related approaches have been used to derive effective low-dimensional models for oscillations induced by chain delays [26–29], and in cases where transcription oscillates stochastically between on and off states [30].

The theory of stochastic systems with random delays is well-understood [31, 32]. The Gillespie algorithm, Langevin equations, and mean-field models can be extended to systems with distributed delays, allowing for efficient sampling at small, intermediate, or high molecule counts, respectively [31, 33, 34]. However, using such models involves a compromise: Adding delay can lead to other challenges, as the resulting models are typically non-Markovian. In particular, this complicates the derivation of parameter likelihood functions, making such models more difficult to analyze and use for parameter inference [35, 36]. Thus, developing a general inference framework for biochemical reaction networks with distributed delays that works even when molecule counts are low remains a challenging open problem. Important progress has been made for certain delay stochastic differential equations [35], and delay linear noise approximations [36]. These approaches rest on the assumption that molecule counts are high enough to allow delay stochastic differential equations to accurately capture system dynamics [31]. However, one must frequently deal with low molecule counts when using imaging data obtained from fluorescent imaging of single cells.

To address this problem, we describe a systematic way to derive likelihoods for the parameters in common biochemical reaction models that include delays. These results allow us to extend MCMC methods to efficiently estimate reaction rates, and delay distribution parameters from single-cell assays even when molecular counts are low. We illustrate the advantages and limitations of our approach using a delay birth/death process, which provides a simple model of gene expression. Our method is robust: It allows us to recover the mean delay even when the delay distribution is misspecified. When only a relative measure of molecule count is available, such as a fluorescence trace, delay parameters can be accurately estimated if the dilution rate can be estimated separately. Our method performs well on experimental data: We show that given the dilution rate of YFP (yellow fluorescent protein) estimated directly from observed cell growth rates, we can infer the time delay of the synthesis of YFP from multiple cell trajectories measured using quantitative time-lapse microscopy. Our approach therefore provides a robust basis for the development of hierarchical networks inference methods that can be used to characterize biochemical processes across cellular populations.

## 2 Methods

### 2.1 Derivation of the likelihood function

Following [6] we consider a chemical reaction network consisting of *u* species, *Y*_1_,…, *Y_u_*, and *v* reactions, *R*_1_,…, *R_v_*. Reaction *R_k_* is given by 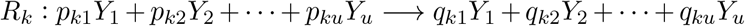, where the *p_kj_* and *q_kj_* are the stoichiometric coefficients. Reaction *R_k_* is equipped with rate constant *θ_k_* and reaction propensity *h_k_*(*y, θ_k_*), where *y*(*t*) = (*y*_1_(*t*), *y*_2_(*t*),…, *y_u_*(*t*)) represents the number of molecules of each chemical species at time *t*. We also define 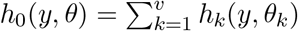, where *θ* = (*θ*_1_, *θ*_2_,…, *θ_v_*).

First assume that we fully observe the state of the chemical reaction network for all *t* ∈ [0, *T*]. Let *r_ki_* denote the number of reactions of type *k* that occur over the time interval (*i, i* + 1], and let 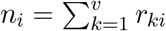. For 1 ⩽ *j* ⩽ *n_i_*, denote the *j*^th^ reaction that occurs within (*i, i* + 1] using the pair (*t_ij_*, *k_ij_*), where reaction *R_k_ij__* occurs at time *t_ij_*.

Suppose that at least one reaction *R_k*_*, once initiated, requires a random time to complete. If the reaction begins at time *t*_initial_, then the time *t*_final_ at which the reaction changes the state of the system is a random variable. We call *t*_final_ – *t*_initial_ the (random) delay associated with reaction *R_k*_*. For instance, production of a given protein starts with the initiation of transcription, but the number of mature proteins in the system changes only after transcription, translation, and post-translational steps result in a fully functional protein.

Let *η_k_* be the measure supported on [0, ∞) that describes the delay distribution associated with reaction *R_k_*. We assume, for the sake of simplicity, that these distributions do not depend on time or the state of the system, and that each *η_k_* depends on a vector of parameters Δ_*k*_ = (Δ_*k*1_, Δ_*k*2_,…, Δ_*kl_k_*_). [33] have proven the existence of reaction *completion* propensities defined by

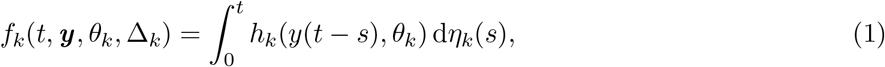

where ***y*** denotes the trajectory of the chemical reaction network from time 0 to time *T*. These propensities define the effective rates of reactions at time *t*, and allow us to write the likelihood of the parameters for an observed sequence of completed reactions in a form analogous to the case without delays [6]. Integrating the completion propensities in time, define

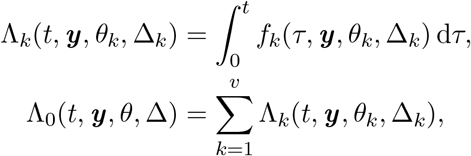

where Δ = {Δ_*kl*_} is the collection of parameters that define all of the delay measures, *η_k_*. If the state, *y*(*t*), of the chemical reaction network is known for all *t* ∈ [0, *T*], then the likelihood function for the set of delay parameters Δ and the vector of rate constants *θ* is given by

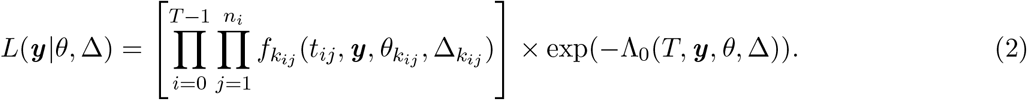

Details of the derivation are provided in the Supplementary Methods. We obtained this likelihood function by adopting a ‘backward’ view of delayed chemical kinetics: We assume that we know *only* the reaction completion times, and treat the corresponding unobserved reaction initiation times that occurred in the past as random quantities. This backward view is useful for the inference problem because reaction initiation times are typically not observed experimentally. By contrast, sampling algorithms such as the Gillespie algorithm typically adopt a ‘forward’ approach, where once a reaction is initiated and recorded, the corresponding reaction completion time is random.

### 2.2 Approximate likelihood given observations at discrete times

Assume now that we only observe the system, *y*(*t*), at a discrete set of times, *t* = 0,1,…, *T* – 1, *T*, yielding a vector of measurements ***y**_d_* = (*y*(0), *y*(1),…, *y*(*T* – 1), *y*(*T*)). These observations can be used to approximate the exact likelihood given by Eq. (2) using a *τ*-leaping approach [32]. First we replace the propensities, *f_k_*, defined in Eq. (1) with propensities, 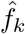, that are constant between observations. We obtain 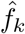 by averaging *f_k_* over [*i, i* + 1], and interpolating *h_k_* linearly between measurements (see Supplementary Methods for details): 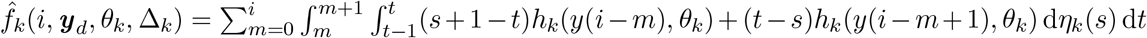, where 0 ⩽ *i* ⩽ *T* – 1, and Δ_*k*_ denotes the vector of parameters that define the measure *η_k_* as before. Note that we do not discretize the delay distributions. Our formula for 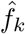 is valid whenever the delay measure *η_k_* is defined by a probability density function. For reactions that do not involve delay (that is, when *η_k_* is a Dirac-delta measure at zero), our formula for 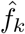 reduces to 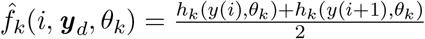.

Using the 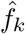 the likelihood in Eq. (2) can be approximated by [31]:

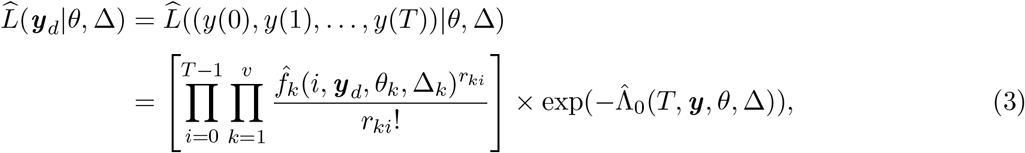

where 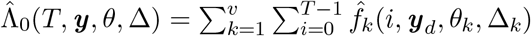, and *r_ki_* denotes the number of reactions of type *k* whose completion has been observed in the interval (*i*, *i* + 1]. When the approximation in Eq. (3) is valid (for instance when the reaction rates are constant between observations), then conditioned on system history up to time *i*, the number of reactions of type *k* that occur within (*i, i* + 1] follows a Poisson distribution with mean 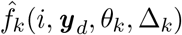.

### 2.3 Statistical inference of model parameters using discrete-time data

We next describe an MCMC algorithm for obtaining the posterior distribution over the model parameters, (*θ*, Δ), using the approximate likelihood given in Eq. (3) and prior distributions over the unknown parameters. Bayes’ Theorem and Eq. (3) allow us to express the posterior distribution over model parameters given the observed data as:

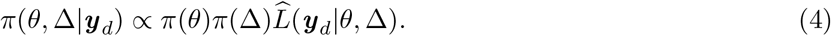

Here *π*(θ) and *π*(Δ) are priors over the rate and delay parameters, respectively. The prior *π*(Δ) can be chosen depending on the delay distributions. We used gamma distributions for *π*(θ) because the support of each *θ_k_* is positive. Moreover, for mass action kinetics the propensity function is separable, so that *h_k_*(*y*(*t*), *θ_k_*) = *θ_k_g_k_*(*y*(*t*)), and hence the gamma distribution defines a conjugate prior [37].

Samples from the posterior distribution given by Eq. (4) can be generated given complete trajectories of all species (i.e. (*t_ij_, k_ij_*)) using Gibbs sampling [15, 38]. In order to sample *θ* and Δ from their conditional posterior distributions, we use the Metroplis-Hastings algorithm [39]. However, sampling *θ* and Δ from their conditional posterior distributions requires knowledge of the number of reactions, *r_ki_*. Crucially, the discrete-time measurements *y*(*i*), *i* = 0,1,…, *T*, do not uniquely determine the number of reactions between observations. We thus need to sample *r_ki_* during each step of Gibbs sampling. To do so, we use the block updating method described by [6] to sample the number of occurrences of each reaction type during each time interval (*i, i* + 1] given the observed system states *y*(*i*) and *y*(*i* + 1), using the Metropolis-Hastings algorithm with a random walk. For the proposal distributions of the number of reactions, we use the Skellam distribution [6, 40]. Since we formulate the posterior in Eq. (4) using an approximate likelihood that reflects a *τ*-leaping approach, we do not consider the specific times at which reactions occur during each time interval (*i, i* + 1], but only their total number (see Supplementary Methods for details).

The following algorithm can then be used to generate samples from the approximate posterior distribution given by Eq. (4).

1. Initialize values for the reaction rates *θ*, parameters for the delay distributions, Δ, and reaction counts *r_ki_*, for the hidden trajectory. Use gamma priors for the rates, *θ_k_*.
2. Sample, in order, *θ_k_*, *k* = 1,…, *v*, from the conditional posterior distribution, given all other rate constants, *θ_ι_*, *l* ≠ *k*, delay parameters Δ, and reaction numbers. If *y*(*t*) and *θ_k_* are separable in the propensity function *h_k_*(*y*(*t*), *θ_k_*), then sample *θ_k_* from the gamma posterior distribution. Otherwise, use the Metropolis-Hastings algorithm.
3. Sample, in order, Δ_*kl*_, 1 ⩽ *k* ⩽ *v* and 1 ⩽ *l* ⩽ *l_k_*, from the conditional posterior, given *θ* and Δ_*k′ι′*_ for all (*k′, l′*) = (*k, l*), using the Metropolis-Hastings algorithm since the conditional posterior does not follow a known distribution.
4. Sample *r_ki_* for all *k* = 1,…,*v* and *i* = 0,…, *T* – 1, given *θ*, Δ, and the observed trajectory, ***y**_d_*, using the block updating method.
5. Repeat steps 2–4 until convergence is achieved.

In the Supplementary Methods we provide all the likelihoods, and illustrate the algorithm in the case of a stochastic birth-death process with delayed birth.

### 2.4 Measurement of YFP level and cell area of individual cells

We tested our algorithm using experimental data from our previous work [41]. Specifically, the P_BAD_-*sfyfp* was cloned to a medium copy-number plasmid, which was later transformed into E. coli JS006A strain which was, in turn, derived from the JS006 strain [42] by introducing a constitutively expressing AraC into the genome. To measure single-cell YFP expression, we cultured the cells in a custom-designed microfluidic device, mounted on a Nikon Eclipse Ti Microscope. Phase contrast and YFP images were taken every 1 minute. Background YFP was first recorded for 12 minutes, then the medium was switched to include 2% ARA to trigger YFP expression. After the experiment, phase-contrast images were segmented and analyzed using custom Matlab code [43]. We analyzed results of two experimental runs. The fluorescence is considerably lower in the second experiment which is likely due to the differences in the heights of the PDMS chip, and conditions of the excitation light bulb between the two experiments.

### 2.5 Relating fluorescence and molecular count

To estimate the fluorescence signal per YFP molecule (*γ*), we measured how the YFP signal of a mother cell is partitioned among two daughter cells. To measure the partitioning during a decreasing phase of the fluoresce signal we followed [44]. We used a genetic circuit with P_BAD_-lacI and Plac/ara-*sfyfp*, which forms an incoherent feedforward loop and thus generates a single pulse of YFP [41]. Specifically, during the decreasing phase after reaching the peak of the pulse, total YFP signals from a mother cell before a cell division and from two daughter cells after the cell division were measured.

## 3 Results

### 3.1 Delay is estimated accurately and precisely with sufficient data

We first tested whether our algorithm can be used to identify the mean, *μ_τ_*, and variance, 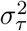, of the delay distribution, as well as the reaction rates of a delayed, stochastic birth-death process:

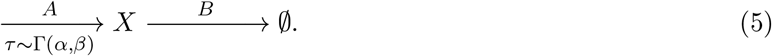

In the generative model, we used gamma distributed delay in the birth reactions, assuming that creation of protein is the result of a chain of exponentially distributed monomolecular reaction steps [19, 20], approximable by a gamma distribution [21, 35, 36]. We generated 500 sample trajectories from the model given by Eq. (5) using the delayed Gillespie algorithm [31], and subsampled each trajectory by recording the molecular count at evenly spaced intervals (Fig. 1a).

**Figure 1:**
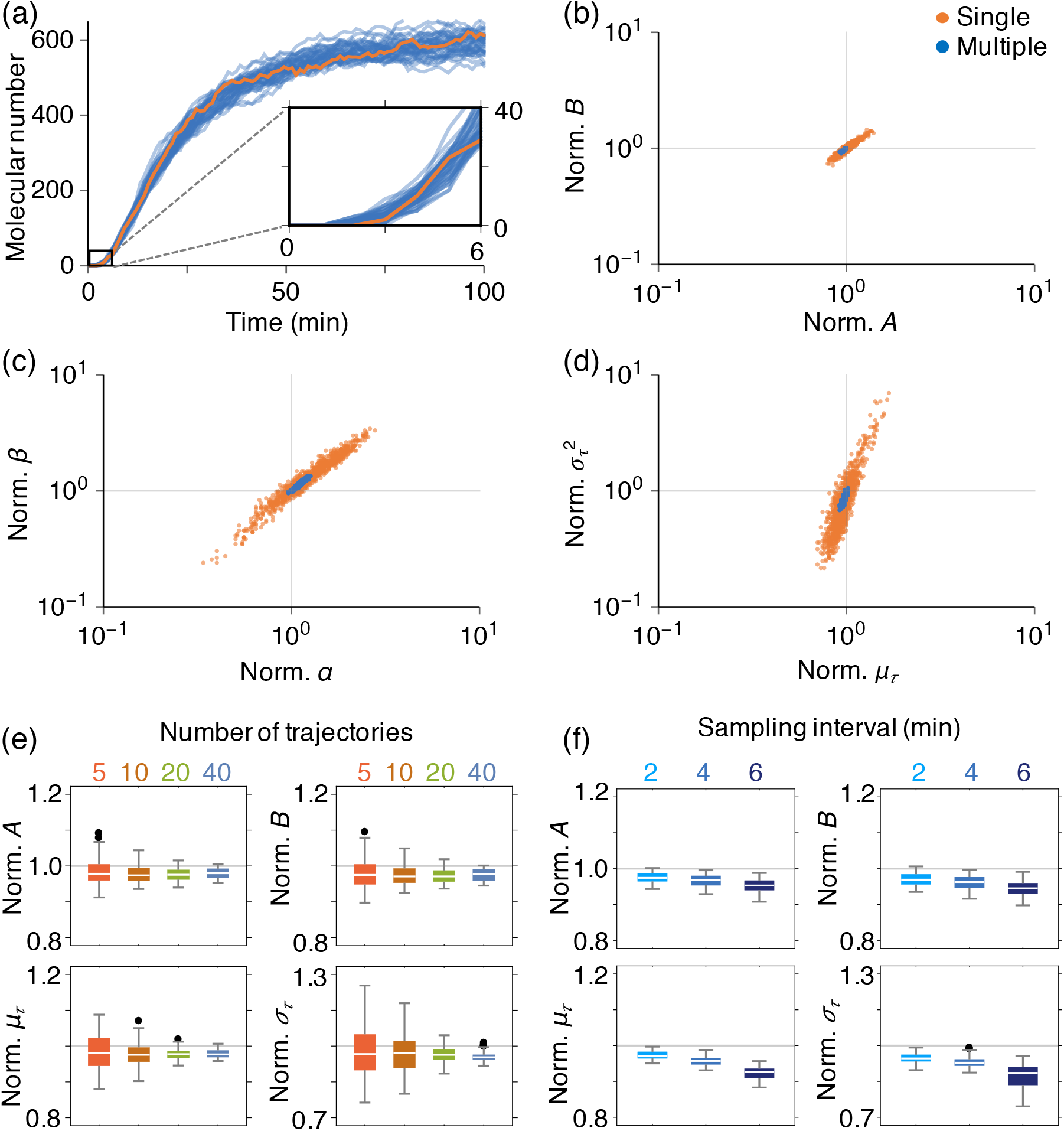
Estimation of the delay distributions using multiple trajectories is accurate and precise. (a) Simulated trajectories of a delayed stochastic birth-death process (Eq. 5) with rate parameters *A* = 30 min^−1^, *B* = 0.05 min^−1^, and delay *τ* ~ Γ(18/5, 5/3), *i.e. μ_τ_* = 6 min and 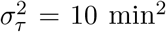. We assumed that *X*(0) = 0. Trajectories used for inference were sampled at I min intervals, (b-d) MCMC generated samples from the posterior distributions over parameters using a single trajectory (orange) or 40 trajectories (blue). While the rate parameters, *A, B*, can be estimated well using a single trajectory (b), estimation of the delay *τ* ~ Γ(*α, β*) requires multiple trajectories (c and d). Here, the sample values were normalized by dividing with the true parameter values, (e) Box plots of 100 posterior means using an increasing number of trajectories. Subsets of between 5 and 40 trajectories were chosen randomly and repeatedly from a set of 500 simulations. The estimates were normalized by dividing with their true parameter values, (f) Sparser measurements at intervals of 2, 4, and 6 min also allowed for accurate and precise estimates of all parameters, as long as a sufficient number of trajectories was used (40 in this case).

A considerable number of molecules was produced before the mean delay time (6 min) due to variability of reaction delays (Fig. 1a inset). Therefore, using either the earliest detectable signal, or a threshold to estimate delay can lead to biased estimates of the mean delay, *μ_τ_*. We inferred the two production and degradation rates, *A, B*, as well as the delay distribution parameters, *α, β*, using the MCMC algorithm described in the Methods and Supplementary Methods. Although we used non-informative priors for all parameters, the reaction rates could be accurately estimated from a single subsampled realization of the process (orange trajectory in Fig. 1a, and posterior distribution in Fig. 1b). However, the posterior distribution over the delay parameters revealed a strong correlation between the two (Fig. 1c), making the delay mean and variance difficult to estimate concurrently (Fig. 1d). Increasing the number of sampled trajectories in the estimation to 40 resulted in an accurate and precise estimate of all parameters (Fig. 1b-d). The accuracy and precision of posterior mean estimates increased with the number of trajectories used for estimation, indicating that the method is consistent (Fig. 1e). Furthermore, with a sufficient number of recordings, even sparsely sampled data still allowed for accurate inference of all model parameters, even when the time between observations equaled the mean delay time (Fig. 1f). These observations are robust for parameters (See Fig. S1). We therefore concluded that our algorithm can be used to estimate delays and rates concurrently, but that multiple trajectories are needed to accurately infer the delay distributions.

### 3.2 Mean delay can be estimated when the underlying time delay distribution is misspecified

We next asked whether delay mean and variance can be estimated even when the delay distribution is misspecified. To do so we generated sample trajectories with delays that followed a beta and inverse gamma distribution with equal mean and variance (See Fig. 2a). While the gamma distribution assumed in the estimation algorithm has infinite support and decays exponentially, the beta distribution has compact support, while the inverse-gamma distribution has a heavy tail.

**Figure 2:**
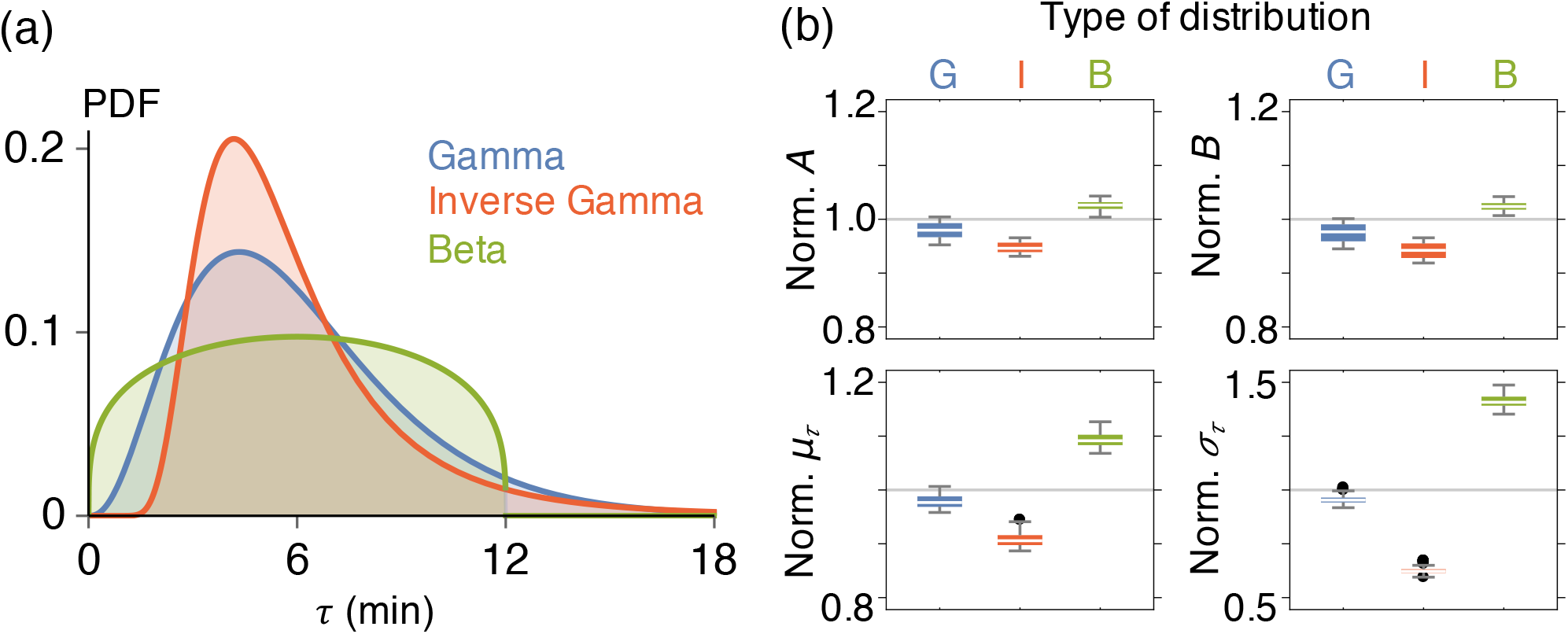
The mean, but not the variance of the delay can be accurately and precisely estimated when the delay distribution is misspecified. (a) Three types of delay distributions: Γ(18/5, 5/3), Inverse-Γ(28/5,138/5), and 12 · *B*(1.3,1.3). For all distributions *μ_τ_* = 6, and 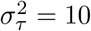. (b) A box plot of 100 posterior mean estimates. As in Fig. 1, parameters were estimated using 40 sample trajectories randomly and repeatedly chosen from a set of 500 trajectories generated assuming one of the thee delay distributions shown in panel (a). Here the estimates were normalized by dividing with their true values.

Given sufficiently many observed trajectories, our algorithm provided accurate and precise estimates of the rates, and mean delay (Fig. 2b). However, the estimate of delay variance, 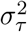, was biased when the delay distribution was misspecified, with a systematic underestimate when the true delay followed an inverse-gamma distribution, and overestimate when the true delay followed a beta distribution.

### 3.3 With relative molecular level measurements estimation of delay requires separate estimates of dilution rate

Frequently we cannot measure the actual molecular count within a cell directly. For instance, measurements of fluorescence reporter intensity are approximately proportional to the absolute species number, but the proportionality constant cannot always be determined precisely, and hence estimates of molecular number from such measurements can be noisy [1, 44–46].

We next asked how such errors in the estimates of absolute protein numbers affect delay distribution inference. To address this question we scaled the sample trajectories in Fig. 1a to mimic a two-fold error in the estimate of the proportionality constant used to convert fluorescence to molecular counts. Such scaling changes the mean and the variance of the signal differently (Fig. 3a) distorting the level of intrinsic fluctuation, as measured by the coefficient of variation. In turn, a mis-scaling can lead to biases in estimation of all parameters including the mean, *μ_τ_*, and variance, 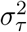, of the delay (Fig. 3b). Thus inaccurate measurements of molecular levels can distort delay estimates.

**Figure 3:**
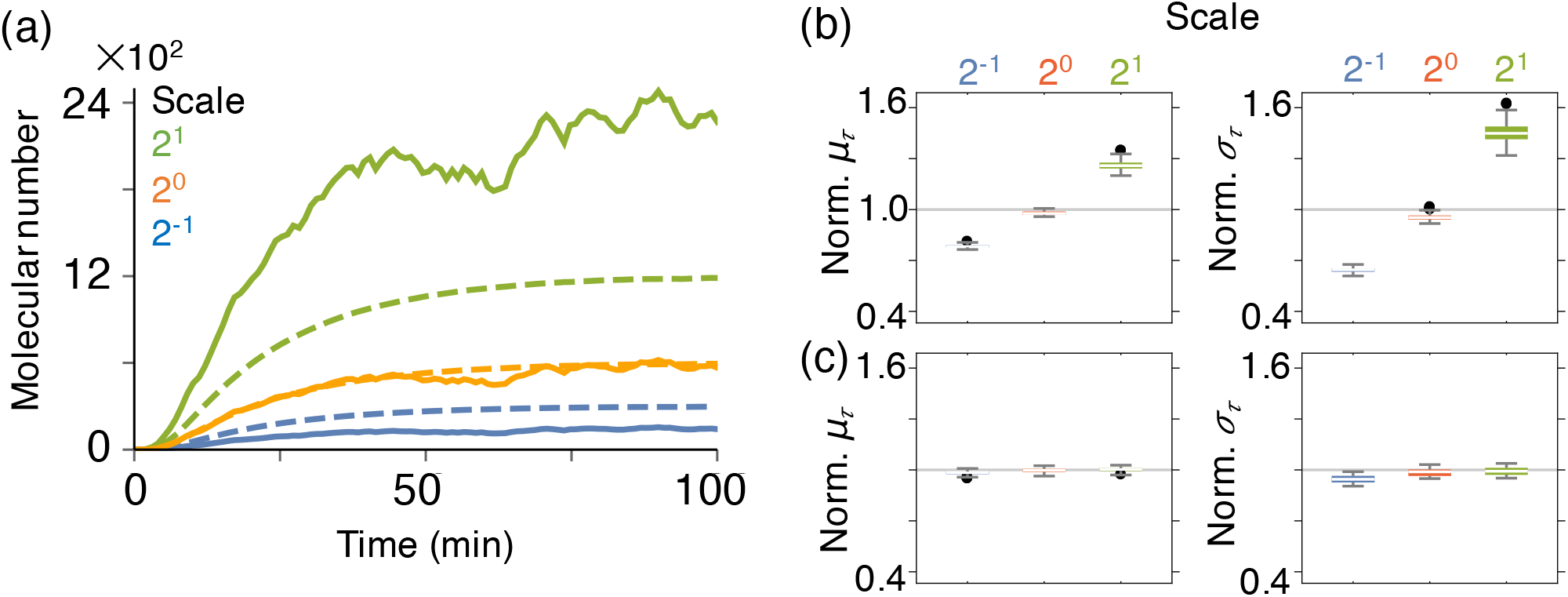
Measurements of dilution rate allow for accurate estimation when only relative molecular levels measurements are available. (a) The average (dashed) and variance (solid) of the 500 simulated trajectories in Fig. 1a, scaled by 0.5, 1, and 2. The average and variance of the scaled trajectories were scaled by different amounts (blue and green). (b) Using scaled trajectories to estimate delays leads to large biases. Here, we show box plots based of 100 posterior means, each estimated using 40 subsampled trajectories. (c) When the dilution rate, *B*, is known, the delay distribution can be accurately and precisely estimated even with incorrectly scaled data.

Advances in lapse imaging techniques are making it easier to estimate cell growth rates, and resulting dilution rates, *B*, directly [23, 47, 48]. We therefore assumed next that the dilution rate, *B*, can be estimated separately, and set to their true value. Once we did so we were able to accurately and precisely estimate delay distribution parameters, even with incorrectly scaled data (Fig. 3c). This indicates that identifying the timescale of the birth-death process by correctly estimating the dilution rate, *B*, can overcome biases due to incorrectly scaled data. Thus our algorithm can be used to effectively infer delays even when only relative molecular levels are known, or when the conversion of fluorescence to protein counts is not accurate.

Furthermore, we found that having access to a separate estimate of the dilution rate, *B*, can resolve unidentifiability issue when only partial data is available. For instance, as their number increases, cells in microfluidic traps can become crowded and their growth can slow as a result [49, 50]. To ensure measurements under minimal strains on the cells, sometimes we use only the initial fluorescence measurements before crowding can impact gene expression (*e.g*. the first 25 min in Fig. 1a). The initial part of the fluorescence trajectory, before saturation, but after YFP maturation, is approximately linear with slope ~ *A/B*. As a consequence only *A/B* is identifiable from data. However, if the dilution rate, *B*, can be estimated separately, the growth rate, *A*, can be estimated even from partial data (Fig. S2). Importantly, the delay parameters, *μ_τ_* and 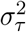, can also be accurately estimated from partial data (Fig. S2). We therefore conclude that a separate measurement of the dilution rate, *B*, allows for successful estimation of time delays from limited, and misscaled data.

### 3.4 Estimation of time delay in transcriptional and translational regulation

We next tested our algorithm on experimental data obtained using time-lapse fluorescence microscopy. As protein synthesis is not instantaneous, there can be considerable delay between gene activation and the formation of functional proteins (Fig. 4a). To estimate this delay, we used a P_BAD_ reporter-only circuit, which we constructed previously by placing a YFP gene under control of the P_BAD_ promoter in *E. coli* [41]. In this circuit the addition of Arabinose (ARA) promotes the rapid activation of AraC, which promotes the constitutive transcription of YFP [23, 24]. Once the translated YFP protein matures, it generates a fluorescence signal. YFP synthesis rate is not strongly affected by cell growth [23, 24, 51]. On the other hand, as cells grow, YFP is diluted. As YFP is a relatively stable protein [52], and is not enzymatically degraded in this system, dilution is the main reason for the decrement in protein number within a cell. Thus, the dynamics of YFP concentration, which is determined by time delayed constitutive synthesis and linear degradation, is well described by Eq. (5).

**Figure 4:**
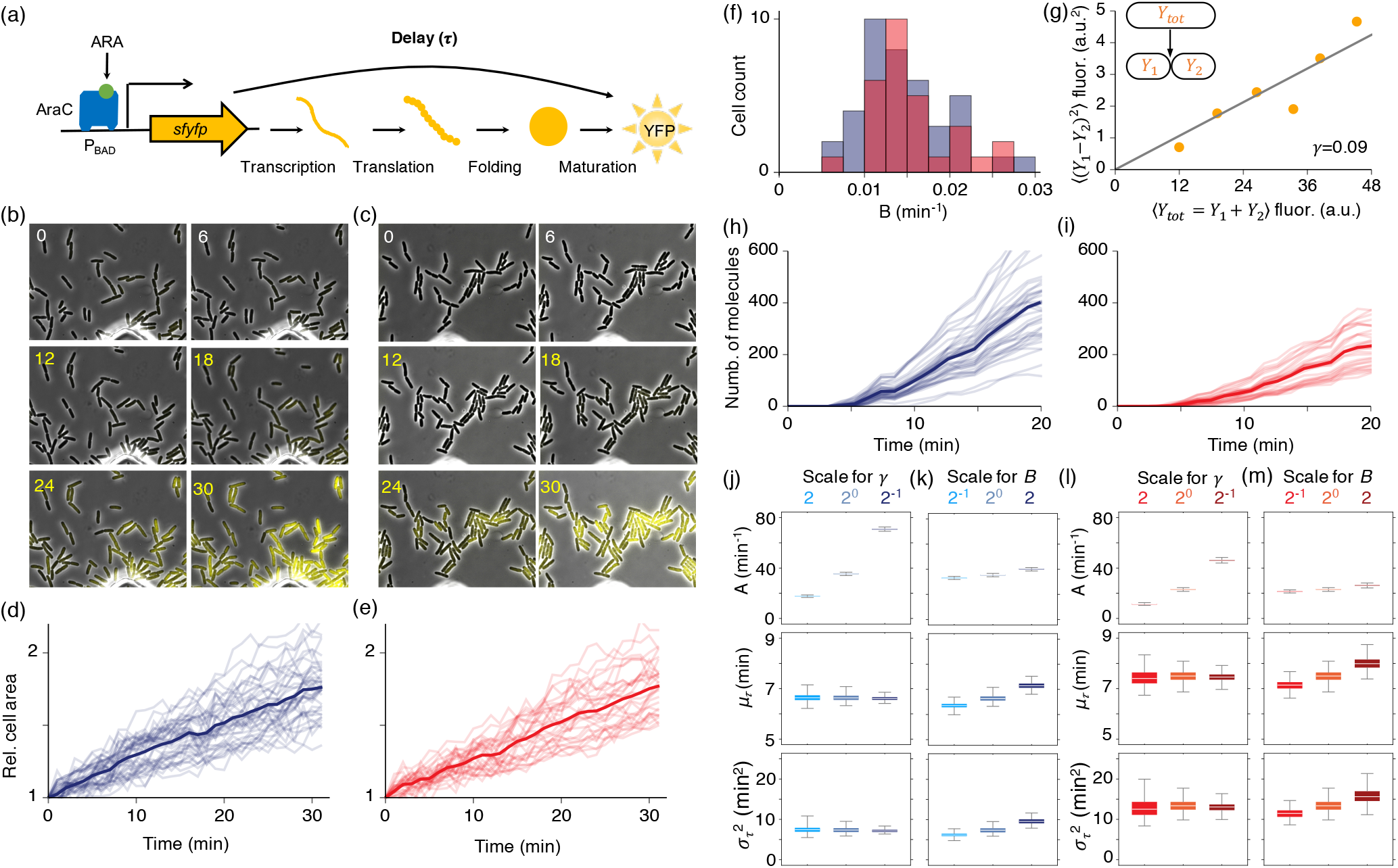
Robust estimation of the time delay distribution of YFP synthesis after induction. (a) When ARA is added to the media, AraC promotes the synthesis of YFP. The synthesis process involves transcription, translation, protein folding and maturation which result in a delay between YFP gene activation, and the observation of the fluorescence signal generated by mature YFP. (b-c) Time-lapse images of YFP expression from two independent experiments, performed previously [41]. At 12 min after measurement was started, 2% ARA was added to the media, promoting the constitutive transcription of YFP. (d-e) The lineage of each cell was identified via manual segmentation of images, and the change in individual cell areas was tracked (39 cells in (d) and 29 cells in (e)). When a mother cell divided into two daughter cells, the area of the mother cell was added to the area of the daughter cell, (f) We estimated dilution rates by fitting an exponential function to the cell growth data. The average dilution rate of the 39 cells from the first, and the 29 cells from the second experiment are 0.015 ± 0.005 and 0.016 ± 0.005 min^−1^, respectively, (g) A conversion constant from YFP signal level to the number of YFP molecules (*γ*) was estimated by measuring the binomial error in partition of total YFP signal (*Y*_tot_) at cell division to two daughther cells (*Y*_1_ and *Y*_2_). The constant *γ* was estimated as described in the text, (h-i) We estimated YFP molecule number per unit area by dividing the total fluorescence level of each individual cell (b, c) with its total area (d, e), and with the estimated scaling factor *γ* (g). (j-m) Using our inference algorithm with these trajectories, and fixing the dilution rate at the estimated value, *B* = 0.015, we obtained 10^4^ posterior samples for the remaining parameters (j and 1). Due to the higher molecular numbers in (h) than (i), the estimated birth rate, *A*, and delay variance, 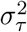, were higher and lower, respectively, in (j) than (1): 35.4 ± 0.4 and 23.1 ± 0.5, and 7.4 ± 0.7 and 13.4 ± 1.4. However, the estimated mean delay time, *μ_τ_*, was similar in the two cases: 6.6 ± 0.1 min (j) and 7.5 ± 0.2 min (1). Estimation of the delay mean and variance was robust to the two-fold change in *γ* (j and 1) and *B* (k and m).

Previously, we performed, and reported on, two independent experiments using time-lapse fluorescence microscopy to measure the YFP trajectories from individual cells in a growing population after induction (Fig. 4b and c) [41]. In both experiments, the fluorescence signal from matured YFP was recorded from each cell at 1 min intervals. After measuring background fluorescence levels for 12 min, we added 2% ARA to the media to promote YFP synthesis (Fig. 4b and c). We tracked the total fluoresce signal in each cell (Fig. 4b and c), to obtain a timeseries of YFP molecule number within a unit area. To do so we first tracked changes in area of individual cells using time–lapse images (Fig. 4d and e). When a cell divided, the area of a the mother cell was added to the area of a daughter cell. By fitting the observed volume growth trajectories to an exponential function, we estimated the dilution rates of individual cells in the population (Fig. 4f), which were consistent with previous estimates [23].

Next, we estimated the fluorescence signal per YFP molecule (*γ*) by estimating the ratio between the square difference of measured fluorescence between two daughter cells ((*Y*_1_ – *Y*_2_)^2^) and the measured fluorescence of a mother cell (*Y*_tot_ = *Y*_1_ + *Y*_2_) (Fig. 4g). This approach is based on the assumption that proteins from the mother cell are partitioned independently, and without bias between the two daughter cells (See [44, 46] for details). Then, by dividing the intensity of the total fluoresce signal in each cell (Fig. 4b and c) by our estimate of *γ*, and the cell’s area (Fig. 4d and e), we obtained an estimate of the timecourse of YFP molecules per unit area for each of 39 cells in the first, (Fig. 4h) and 29 cells in the second experiment (Fig. 4i).

In neither experiment did the YFP signal saturate before the end of the experiment, and we thus only obtained partial trajectories in both cases. To address this problem, we fixed the dilution rate, *B* = 0.015 (Fig. 4f) which we estimated from the observed rate of growth and division while estimating the remaining parameters, *A, μ_τ_* and 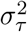 from the partial trajectories. The difference in the total fluorescence levels between the two experiments (Fig. 4h and i) resulted in a higher estimated production rate, *A*, in the first experiment (Fig. 4j). Without investigating further, we could not tell whether this difference in production rates was real, or whether discrepancies in the experimental setup caused a difference in the strengths of the recorded signal.

Despite the difference in the inferred rates, the estimated mean delay times, *μ_τ_*, were similar: 6.6 ± 0.1 min (Fig. 4j) and 7.5 ± 0.2 min (Fig. 4l), in the first and second experiment, respectively. The estimated time delay is similar to the time to maturation of YFP variant VENUS (7 ± 2.5) measured using real-time monitoring of a single molecule [45], supporting the accuracy of our algorithm.

The inferred proportionality constant, *γ* = 0.09, (Fig. 4g) depends on the camera setting, and can vary between experiments (Fig. 4h and i). We thus examined the impact of varying the constant *γ* on the estimated parameters. Even in the presence of a two-fold change in *γ*, the estimates of delay mean, *μ_τ_*, and variance, 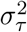, changed little (Fig. 4j and l). Thus our conclusions about the robustness of the inference algorithm when the dilution rate is known extend to experimental data (Fig. 3c). Furthermore, as the dilution rate, *B*, can differ between cells (Fig. 4f), we also investigated the sensitivity of our inference method to changes in the exact value of this constant. Even a two-fold change in the dilution, resulted in only a small changes in the estimate of mean delay time, *μ_τ_* (~ 5%; See Fig. 4k and m), providing a further indication that our approach is robust.

## 4 Conclusion

We have introduced a principled approach to extending Bayesian inference techniques that allows for parameter estimation in biochemical reaction networks with delays. We have shown that the method can be used to estimate both reaction rates and delay distribution parameters from experimentally-obtainable observation of gene regulatory networks. Although the method has some limitations, we have shown that they can be addressed by proper experimental design.

We considered a simple birth-death process with a small number of parameters in order to understand the advantages and limitations of the proposed method. Nevertheless, our approach is scalable: The derivation of the likelihood function for the different parameters, and the experimental design principles we discussed can be extended to systems with many biochemical species, multiple delays, and complex dynamics. Examples include networks of interacting birth-death processes with nonlinear delayed protein synthesis, and systems that oscillate due to delayed negative feedback loops [28, 41]. Importantly, replacing unobserved or uninteresting reaction pathways with time delays in large biochemical reaction networks can significantly reduce the number of model parameters. We thus expect that an equivalent algorithm to the one we presented can then be used to infer rates and characterize delays in the resulting reduced networks. The identifiability of time delay in more complex models is a challenge that we will address in future work.

When molecular counts are sufficiently high, chemical master equations can be approximated by analytically tractable reductions such as delay stochastic differential equation (SDEs), and linear noise approximations (LNAs) [31, 34, 53, 54]. Previous work has leveraged these approximations for Bayesian parameter inference. Specifically, [35] have developed a Bayesian algorithm using SDE models containing distributed delay, with particular emphasis on oscillations generated by delayed negative feedback loops [55]. Recently, a filtering approach based on LNAs has been developed to infer distributed delays [36]. An interesting avenue for future research is to develop hybrid models, and combine our method with previous SDE or LNA approaches to gain both in computational speed and accuracy.

While delay distributions were difficult to infer from a single trajectory, a relatively small number of trajectories allowed for efficient inference of all parameters. An important caveat is that when we used multiple cell trajectories for inference, we assumed that all recorded cells were identical. Thus, our algorithm at present does not take into account cell-to-cell variability in YFP expression due to differences in growth rates, plasmid copy numbers, asymmetric partition of proteins at division, and other factors.

In particular, heterogeneity in ARA uptake rates is known to cause considerable cell-to-cell variation in time delay [23]. However, the 2% ARA in the media we used in our experiments was sufficiently high to ensure that uptake occurred rapidly, and minimized cell-to-cell variability. This at least partly justifies our assumption that cell-to-cell differences in time delay are mainly due to measurement, and intrinsic noise. Indeed, our estimates of time delay are consistent with those obtained using real-time monitoring of a single molecule [45]. The robust performance of our method with relatively small number of measurements suggests that it can be extended to hierarchical models which take into account cell-to-cell variability and extrinsic noise sources [9].

In sum, we have presented a method to characterize reduced models of biochemical networks with delays. Our approach is flexible, and the robustness of the method suggests that it can be extended to more complex biochemical reaction networks, and hierarchical models allowing us to shine a light on complex processes within cells, and populations.

## Acknowledgements

We thank Alan Veliz-Cuba and Mehdi Sadeghpour for valuable comments.

## References

1. Long Cai, Nir Friedman, and X Sunney Xie. Stochastic protein expression in individual cells at the single molecule level. Nature, 440(7082):358, 2006.

2. Mads Kaern, Timothy C Elston, William J Blake, and James J Collins. Stochasticity in gene expression: from theories to phenotypes. Nature Reviews Genetics, 6(6):451, 2005.

3. Thomas B Kepler and Timothy C Elston. Stochasticity in transcriptional regulation: origins, consequences, and mathematical representations. Biophysical journal, 81(6):3116–3136, 2001.

4. Arjun Raj and Alexander van Oudenaarden. Nature, nurture, or chance: stochastic gene expression and its consequences. Cell, 135(2):216–226, 2008.

5. Stephen Smith and Ramon Grima. Single-cell variability in multicellular organisms. Nature Communications, 9(1):345, 2018.

6. Richard J Boys, Darren J Wilkinson, and Thomas BL Kirkwood. Bayesian inference for a discretely observed stochastic kinetic model. Statistics and Computing, 18(2):125–135, 2008.

7. Suresh Kumar Poovathingal and Rudiyanto Gunawan. Global parameter estimation methods for stochastic biochemical systems. BMC bioinformatics, 11(1):414, 2010.

8. Bernie J Daigle, Min K Roh, Linda R Petzold, and Jarad Niemi. Accelerated maximum likelihood parameter estimation for stochastic biochemical systems. BMC bioinformatics, 13(1):68, 2012.

9. Christoph Zechner, Michael Unger, Serge Pelet, Matthias Peter, and Heinz Koeppl. Scalable inference of heterogeneous reaction kinetics from pooled single-cell recordings. Nature methods, 11(2):197, 2014.

10. Bernie J Daigle Jr, Mohammad Soltani, Linda R Petzold, and Abhyudai Singh. Inferring single-cell gene expression mechanisms using stochastic simulation. Bioinformatics, 31(9):1428–1435, 2015.

11. Christoph Zimmer, Sven Sahle, and Jürgen Pahle. Exploiting intrinsic fluctuations to identify model parameters. IET systems biology, 9(2):64–73, 2015.

12. Frank T Bergmann, Sven Sahle, and Christoph Zimmer. Piecewise parameter estimation for stochastic models in copasi. Bioinformatics, 32(10):1586–1588, 2016.

13. Boseung Choi, Grzegorz A Rempala, and Jae Kyoung Kim. Beyond the michaelis-menten equation: Accurate and efficient estimation of enzyme kinetic parameters. Scientific reports, 7(1):17018, 2017.

14. Andrew Golightly and Darren J Wilkinson. Bayesian inference for stochastic kinetic models using a diffusion approximation. Biometrics, 61(3):781–788, 2005.

15. Boseung Choi and Grzegorz A Rempala. Inference for discretely observed stochastic kinetic networks with applications to epidemic modeling. Biostatistics, 13(1):153–165, 2012.

16. Michael GB Blum, Maria Antonieta Nunes, Dennis Prangle, Scott A Sisson, et al. A comparative review of dimension reduction methods in approximate bayesian computation. Statistical Science, 28(2):189–208, 2013.

17. Marcella M Gomez, Richard M Murray, and Matthew R Bennett. The effects of time-varying temperature on delays in genetic networks. SIAM journal on applied dynamical systems, 15(3):1734–1752, 2016.

18. Anja Korenčič, Grigory Bordyugov, Damjana Rozman, Marko Goličnik, Hanspeter Herzel, et al. The interplay of cis-regulatory elements rules circadian rhythms in mouse liver. PloS one, 7(11):e46835, 2012.

19. Manuel Barrio, André Leier, and Tatiana T Marquez-Lago. Reduction of chemical reaction networks through delay distributions. The Journal of chemical physics, 138(10):104114, 2013.

20. Andre Leier, Manuel Barrio, and Tatiana T Marquez-Lago. Exact model reduction with delays: closed-form distributions and extensions to fully bi-directional monomolecular reactions. Journal of The Royal Society Interface, 11(95):20140108, 2014.

21. Golan Bel, Brian Munsky, and Ilya Nemenman. The simplicity of completion time distributions for common complex biochemical processes. Physical biology, 7(1):016003, 2009.

22. Ido Golding, Johan Paulsson, Scott M Zawilski, and Edward C Cox. Real-time kinetics of gene activity in individual bacteria. Cell, 123(6):1025–1036, 2005.

23. Judith A Megerle, Georg Fritz, Ulrich Gerland, Kirsten Jung, and Joachim O Rädler. Timing and dynamics of single cell gene expression in the arabinose utilization system. Biophysical journal, 95(4):2103–2115, 2008.

24. Georg Fritz, Judith A Megerle, Sonja A Westermayer, Delia Brick, Ralf Heermann, Kirsten Jung, Joachim O Rädler, and Ulrich Gerland. Single cell kinetics of phenotypic switching in the arabinose utilization system of e. coli. PLoS One, 9(2):e89532, 2014.

25. Harley H McAdams and Lucy Shapiro. Circuit simulation of genetic networks. Science, 269(5224):650–656, 1995.

26. William Mather, Matthew R Bennett, Jeff Hasty, and Lev S Tsimring. Delay-induced degrade-and-fire oscillations in small genetic circuits. Physical review letters, 102(6):068105, 2009.

27. Manuel Barrio, Kevin Burrage, André Leier, and Tianhai Tian. Oscillatory regulation of hes1: discrete stochastic delay modelling and simulation. PLoS computational biology, 2(9):e117, 2006.

28. Ye Chen, Jae Kyoung Kim, Andrew J Hirning, Krešimir Josić, and Matthew R Bennett. Emergent genetic oscillations in a synthetic microbial consortium. Science, 349(6251):986–989, 2015.

29. Faiza Hussain, Chinmaya Gupta, Andrew J Hirning, William Ott, Kathleen S Matthews, Krešimir Josić, and Matthew R Bennett. Engineered temperature compensation in a synthetic genetic clock. Proceedings of the National Academy of Sciences, 111(3):972–977, 2014.

30. Julian Lewis. Autoinhibition with transcriptional delay: a simple mechanism for the zebrafish somitogenesis oscillator. Current Biology, 13(16):1398–1408, 2003.

31. Chinmaya Gupta, José Manuel López, Robert Azencott, Matthew R Bennett, Krešimir Josić, and William Ott. Modeling delay in genetic networks: From delay birth-death processes to delay stochastic differential equations. The Journal of chemical physics, 140(20):05B624_1, 2014.

32. Ankur Gupta and James B Rawlings. Comparison of parameter estimation methods in stochastic chemical kinetic models: Examples in systems biology. AIChE Journal, 60(4):1253–1268, 2014.

33. Robert Schlicht and Gerhard Winkler. A delay stochastic process with applications in molecular biology. Journal of mathematical biology, 57(5):613–648, 2008.

34. Tobias Brett and Tobias Galla. Stochastic processes with distributed delays: Chemical Langevin equation and linear noise approximation. Physical Review Letters, 110(25):250601, 2013.

35. Elizabeth A Heron, Bärbel Finkenstadt, and David A Rand. Bayesian inference for dynamic transcriptional regulation; the hes1 system as a case study. Bioinformatics, 23(19):2596–2603, 2007.

36. Silvia Calderazzo, Marco Brancaccio, and Bärbel Finkenstädt. Filtering and inference for stochastic oscillators with distributed delays. Bioinformatics, 2018.

37. Darren J Wilkinson. Stochastic Modelling for Systems Biology. CRC Press, 2nd. edition, 2011.

38. Adrian FM Smith and Gareth O Roberts. Bayesian computation via the gibbs sampler and related markov chain monte carlo methods. Journal of the Royal Statistical Society. Series B (Methodological), pages 3–23, 1993.

39. Luke Tierney. Markov chains for exploring posterior distributions. the Annals of Statistics, page 1701–1728, 1994.

40. Norman Johnson and Samuel I Kotz. Distributions in Statistics: Discrete distributions V. 3. John Wiley & Sons, 1985.

41. Yu-Yu Cheng, Andrew J Hirning, Krešimir Josić, and Matthew R Bennett. The timing of transcriptional regulation in synthetic gene circuits. ACS synthetic biology, 6(11):1996–2002, 2017.

42. Jesse Stricker, Scott Cookson, Matthew R Bennett, William H Mather, Lev S Tsimring, and Jeff Hasty. A fast, robust and tunable synthetic gene oscillator. Nature, 456(7221):516, 2008.

43. Alan Veliz-Cuba. https://github.com/alanavc/rodtracker, 2014.

44. Nitzan Rosenfeld, Jonathan W Young, Uri Alon, Peter S Swain, and Michael B Elowitz. Gene regulation at the single-cell level. science, 307(5717):1962–1965, 2005.

45. Ji Yu, Jie Xiao, Xiaojia Ren, Kaiqin Lao, and X Sunney Xie. Probing gene expression in live cells, one protein molecule at a time. Science, 311(5767):1600–1603, 2006.

46. Nitzan Rosenfeld, Theodore J Perkins, Uri Alon, Michael B Elowitz, and Peter S Swain. A fluctuation method to quantify in vivo fluorescence data. Biophysical journal, 91(2):759–766, 2006.

47. Thomas M Norman, Nathan D Lord, Johan Paulsson, and Richard Losick. Memory and modularity in cell-fate decision making. Nature, 503(7477):481, 2013.

48. Sattar Taheri-Araghi, Serena Bradde, John T Sauls, Norbert S Hill, Petra Anne Levin, Johan Paulsson, Massimo Vergassola, and Suckjoon Jun. Cell-size control and homeostasis in bacteria. Current Biology, 25(3):385–391, 2015.

49. Dmitri Volfson, Scott Cookson, Jeff Hasty, and Lev S Tsimring. Biomechanical ordering of dense cell populations. Proceedings of the National Academy of Sciences, 105(40):15346–15351, 2008.

50. Morgan Delarue, Jörn Hartung, Carl Schreck, Pawel Gniewek, Lucy Hu, Stephan Herminghaus, and Oskar Hallatschek. Self-driven jamming in growing microbial populations. Nature physics, 12(8):762, 2016.

51. DW Austin, MS Allen, JM McCollum, RD Dar, JR Wilgus, GS Sayler, NF Samatova, CD Cox, and ML Simpson. Gene network shaping of inherent noise spectra. Nature, 439(7076):608, 2006.

52. Jens Bo Andersen, Claus Sternberg, Lars Kongsbak Poulsen, Sara Petersen Bjørn, Michael Givskov, and Søren Molin. New unstable variants of green fluorescent protein for studies of transient gene expression in bacteria. Applied and environmental microbiology, 64(6):2240–2246, 1998.

53. Jae Kyoung Kim, Krešimir Josić, and Matthew R Bennett. The validity of quasi-steady-state approximations in discrete stochastic simulations. Biophysical journal, 107(3):783–793, 2014.

54. P. Thomas, A. V. Straube, and R. Grima. The slow-scale linear noise approximation: an accurate, reduced stochastic description of biochemical networks under timescale separation conditions. BMC Syst. Biol., 6, 2012.

55. Nicholas AM Monk. Oscillatory expression of hes1, p53, and nf-*κ*b driven by transcriptional time delays. Current Biology, 13(16):1409–1413, 2003.

